# The opportunistic intracellular bacterial pathogen *Rhodococcus equi* elicits type I interferons by engaging cytosolic DNA sensing in macrophages

**DOI:** 10.1101/2021.03.28.437424

**Authors:** Krystal J Vail, Bibiana Petri da Silveira, Samantha L Bell, Angela I Bordin, Noah D Cohen, Krisitn L Patrick, Robert O Watson

## Abstract

*Rhodococcus equi* is a major cause of foal pneumonia and an opportunistic pathogen in immunocompromised humans. While alveolar macrophages constitute the primary replicative niche for *R. equi*, little is known about how intracellular *R. equi* is sensed by macrophages. Here, we discovered that that in addition to previously characterized pro-inflammatory cytokines (e.g., Tnfa, Il6, Il1b), macrophages infected with *R. equi* induce a robust type I IFN response, including *Ifnb* and interferon-stimulated genes (ISGs), similar to the evolutionarily related pathogen, *Mycobacterium tuberculosis*. Follow up studies using a combination of mammalian and bacterial genetics, demonstrated that induction of this type I IFN expression program is largely dependent on the cGAS/STING/TBK1 axis of the cytosolic DNA surveillance pathway, suggesting that *R. equi* perturbs the phagosomal membrane and causes DNA release into the cytosol following phagocytosis. Consistent with this we found that a population of ~12% of *R. equi* phagosomes recruited the galectin-3, −8 and −9 danger receptors. Interesting, neither phagosomal damage nor induction of type I IFN required the *R. equi*’s virulence-associated plasmid. Importantly, *R. equi* infection of both mice and foals stimulated ISG expression, in organs (mice) and circulating monocytes (foals). By demonstrating that *R. equi* activates cytosolic DNA sensing in macrophages and elicits type I IFN responses in animal models, our work provides novel insights into how *R. equi* engages the innate immune system and furthers our understanding how this zoonotic pathogen causes inflammation and disease.

**IMPORTANCE:** *Rhodococcus equi* is a facultative intracellular bacterial pathogen of horses and other domestic animals, as well as an opportunistic pathogen of immunocompromised and rarely immunocompetent humans. In human patients, *Rhodococcus* pneumonia bears some pathological similarities to pulmonary tuberculosis, and poses a risk for misdiagnosis. In horses, *R. equi* infection has a major detrimental impact on the equine breeding industry due to a lack of an efficacious vaccine and its ubiquitous distribution in soil. Given the prevalence of subclinical infection and high false positive rate in current screening methods, there exists a critical need to identify factors contributing to positive patient outcomes. Our research identifies innate immune sensing events and immune transcriptional signatures that may lead to biomarkers for clinical disease, more accurate screening methods, and insight into susceptibility to infection.

## INTRODUCTION

*Rhodococcus equi* is a gram positive, intracellular bacterial pathogen that causes severe, potentially fatal respiratory disease in young horses up to 6 months of age. *R. equi* infection has a major detrimental impact on the equine breeding industry for several reasons: it is nearly ubiquitous in the soil of some facilities, there is not currently an efficacious vaccine, and early diagnosis is challenging (1). Virtually all foals are exposed by inhalation of contaminated soil. While many develop subclinical infection, 18-50% of foals develop pneumonia but recover with treatment, and 2-5% perish (2–5). Those that do not succumb to disease develop lifelong immunity and active infection is rare in adult horses (6).

*R. equi* is also a pathogen of humans, causing a pneumonia that radiographically and pathologically resembles pulmonary tuberculosis (TB), as well as extrapulmonary infections (7–9). The majority of human cases manifest as pneumonia and occur in immunocompromised individuals, such as those with impaired cell mediated immunity due to HIV infection (10) or immunosuppression therapy related to organ transplantation (11). However, a growing number of cases have been reported in immunocompetent humans, less than half of which develop pulmonary lesions (8, 9). Over 50% of human infections are derived from porcine- or equine-adapted strains, indicating that most human *R. equi* infections are zoonotic (10, 12). Upon inhalation, *R. equi* survives and replicates within alveolar macrophages in a phagosomal compartment that fails to mature into a lysosome, resulting in the *R. equi*-containing vacuole. Previous studies have shown that *R. equi* pathogenesis depends in large part on expression of bacterial virulence factors (13–16) and production of pro-inflammatory cytokines (17–19), but the nature of the innate immune milieu generated by *R. equi* infection remains ill-defined.

The toll-like receptor (TLR) family is a vital component of innate immunity against microbial pathogens. TLR signaling, through adaptor proteins such as MyD88 and TRIF, as well as transcription factors like NF-κB, is central to pro-inflammatory cytokine production (20). Several lines of evidence demonstrate the importance of host expression of pro-inflammatory cytokines such as IFN-g and TNF-a in *R. equi* pathogenesis. While most laboratory strains of mice are resistant to *R. equi*, treatment with monoclonal antibodies against IFN-g, TNF-a, or both fail to clear *R. equi* infection, develop pulmonary lesions, and succumb to disease (18, 19, 21). Likewise, *R. equi* replication is reduced in equine monocyte derived macrophages primed with IFN-g or TNF-a prior to infection (22). The requirement for pro-inflammatory cytokine production in clearing *R. equi* was further illustrated by Darrah and colleagues, who showed that IFN-g deficient (*Ifng^-/-^*) mice infected with low dose *R. equi* were hypersusceptible to infection (died 13 days post-infection), and mice with impaired nitric oxide (*Nos2^-/-^*) or superoxide (*Gp91^phox-/-^*) production were even more susceptible and died by days 7.5 and 9.5 post infection respectively (23). Together, these studies suggest that IFN-g activates macrophages to produce reactive oxygen species that limit intracellular replication and kill *R. equi* (23).

While signaling via IFN-g, a type II IFN, is important for macrophage activation and control of bacterial infection, type I IFN can act as a negative regulator of host defenses against intracellular bacterial infection. This paradigm is particularly evident in mycobacterial infection, where a type II IFN signature is associated with mild pathogenesis and increased pathogen clearance during both *Mycobacterium tuberculosis* and *Mycobacterium leprae* infection: a type I IFN response is correlated with diffuse lepromatous leprosy and active tuberculosis in humans (24, 25). Both *M. leprae* and *M. tuberculosis* replicate within a modified phagosome, which they permeabilize via their ESX-1 virulence-associated secretion systems, to interact with the host macrophage. It is increasingly appreciated that numerous intracellular bacterial pathogens, including *M. tuberculosis*, *L. monocytogenes*, and *F. tularensis*, activate cytosolic DNA sensing and induce type I IFN signaling through similar mechanisms (26–28). Like mycobacteria, *R. equi* also possesses an ESX secretion system (29), but the contribution of cytosolic surveillance to the macrophage response to *R. equi* has not known.

In spite of the detrimental impact this important pathogen has on the equine breeding industry, data on macrophage sensing of *R. equi* is incomplete, and to date, studies of this pathogen have centered on extracellular macrophage receptors. Here, we sought to investigate whether this vacuolar pathogen triggers innate immune responses in the macrophage cytosol. Additionally, we sought to characterize the transcriptional response triggered by *R. equi* during *ex vivo* infection in murine macrophages as well as *in vivo* in mice and in foals. Using RNA-seq, we found that numerous type I IFN genes were upregulated during *R. equi* infection and that this bacterium induces phagosomal permeabilization in a way that recruits galectin danger sensors. While both galectin recruitment and the type I IFN gene expression profile was not dependent on expression of the *R. equi* virulence-associated protein A (VapA), type I IFN production did require the cGAS/STING/TBK1 signaling axis. Furthermore, we found that a type I IFN program was induced *in vivo* in both a mouse model as well as in an equine model. These data provide evidence that *R. equi* activates the cytosolic DNA sensing pathway during macrophage infection and suggest that type I IFN signaling may be critical for *R. equi* pathogenesis.

## RESULTS

### Transcriptomics uncovers upregulation of pro-inflammatory cytokines and type I IFN in *R. equi*infected murine macrophages

To begin to define the nature of the macrophage innate immune response to *R. equi*, we turned to RNA-seq as an unbiased approach to assess global gene expression changes following infection. Briefly, RAW 264.7 macrophages were infected with virulent *R. equi* (ATCC 33701+) at a multiplicity of infection (MOI) of 5, and total RNA from uninfected and infected cells was harvested after 4h (previous studies of Mtb infection of RAW 264.7 cells show robust induction of TLR and cytosolic nucleic acid sensing pathways at this time point) (28, 30, 31). High-throughput RNA sequencing was carried out on 3 biological replicates of uninfected and infected macrophages. After filtering transcripts with a fold change of > ± 2 (*p* < 0.05), we identified 390 genes that were upregulated and 65 genes that were downregulated in *R. equi* infected macrophages compared to uninfected controls (Fig. 1A). Interestingly, the expression profile had considerable overlap with that of macrophages infected with Mtb, with 167 *R. equi*-induced genes also upregulated in Mtb-infected macrophages, and 6 genes downregulated in both groups (Fig. 1B). Consistent with previous findings (17, 32, 33), we observed significant upregulation of numerous canonical pro-inflammatory cytokines (*Il1a, Il1b*, and *Tnf*), chemokines (*Cxcl2, Ccl4, Ccl3, Cxcl10*), inflammasome genes (*Nlrp3*), and prostaglandins (*Ptgs2*) at 4h post-*R. equi* infection (Fig. 1C). We also observed upregulation of several antiviral genes that are induced downstream of the IRF (interferon regulatory factor) family of transcription factors (*Rsad2*, *Ifit1*). To validate RNA-seq gene expression changes during *R. equi* infection, representative upregulated (*Lif, Nlrp3*) and downregulated (*Mafb* and *S1pr1*) transcripts were measured by RT-qPCR (Fig. 1D). Because some innate immune transcripts can peak after 4h, we measured gene expression at both 4- and 8h following infection; these RT-qPCR results were consistent with the RNA-seq data.

**Figure 1.**
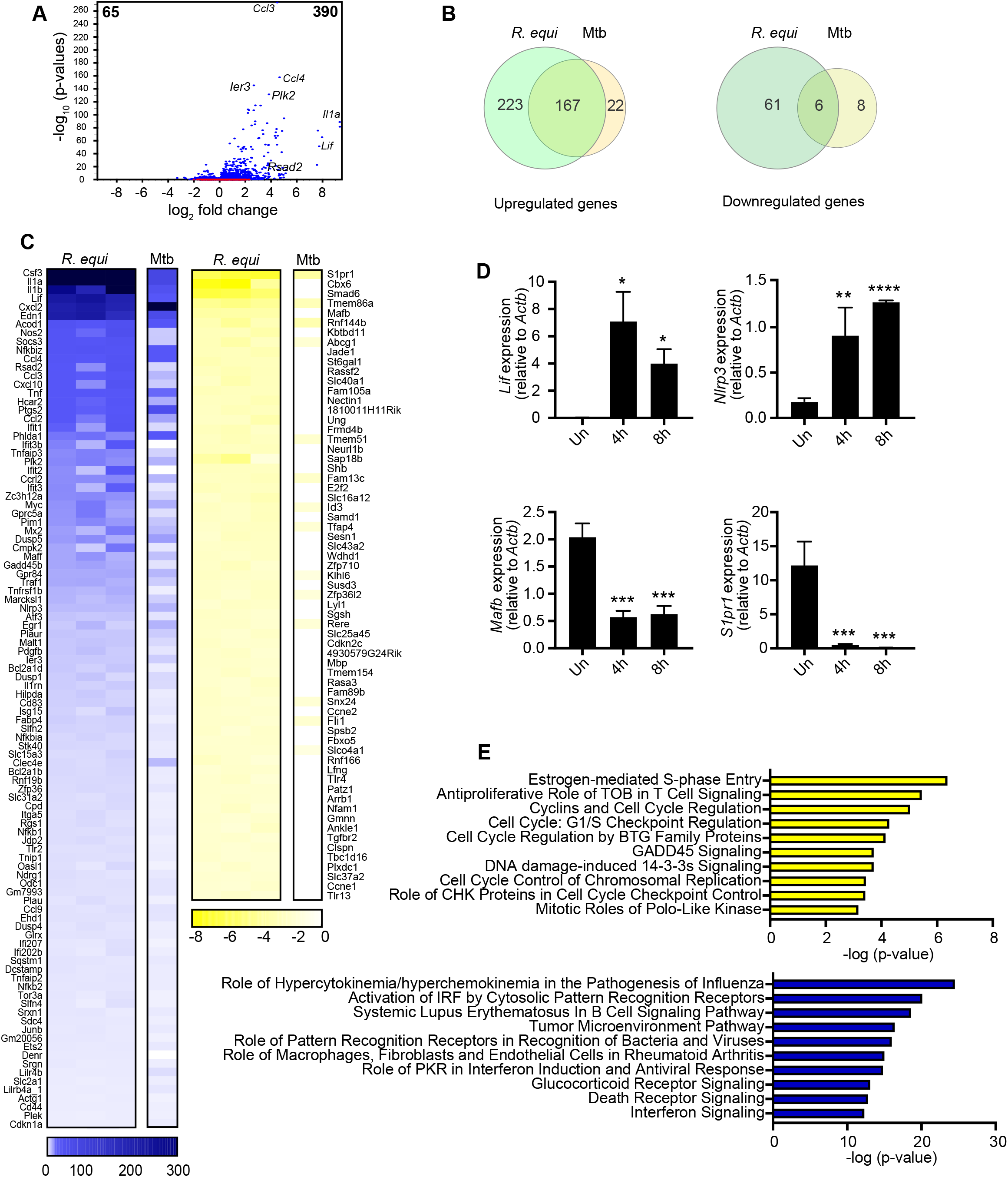
Transcriptomics reveal upregulation of pro-inflammatory and type I IFN response genes in R. equi-infected murine macrophages. (A) Volcano plot of gene expression analysis of R. equi infected macrophages. x axis shows fold change of gene expression and y axis shows statistical significance. Downregulated genes are plotted on the left and upregulated genes are on the right. (B) Venn diagram comparing differentially expressed genes in R. equi- or Mtb-infected RAW 264.7 macrophages. (C) Heatmap showing gene expression analysis of RAW 264.7 macrophages infected for 4 hours with R. equi or Mtb compared to uninfected cells (upregulated genes in blue, downregulated in yellow). Each column for R. equi represents a biological replicate. Mtb is shown as average of 3 replicates. (D) qRT-PCR validation of upregulated genes (Lif, Nlrp3) and downregulated genes (Mafb, S1pr1) in RAW 264.7 macrophages at 4 and 8 hours post infection with R. equi. (E) Ingenuity pathway analysis of gene expression changes in R. equi-infected RAW 264.7 macrophages (upregulated genes in blue, downregulated in yellow). (A-E) represent 3 biological replicates ± SD, n=3. For qRT-PCRs, statistical significance was determined using Students’ t-test. *p < 0.05, **p < 0.01, ***p < 0.001, ****p < 0.0001, ns = not significant. For DGE statistical significance was determined using the EDGE test in CLC Genomics Workbench. Significant differentially expressed genes were those with a p <0.05 and fold change of <±2.

Using Ingenuity Pathway Analysis (IPA, Qiagen), we next asked which pathways were enriched for differentially expressed genes (DEG) in uninfected vs. *R. equi*-infected macrophages. Unbiased canonical pathway analysis of the biological processes most enriched during *R. equi* infection showed strong upregulation of genes related to innate immune signaling (“Role of pattern recognition receptors in recognition of bacteria and viruses Interferon signaling”, “Glucocorticoid receptor signaling”), cell death (“Death receptor signaling”), and tumor pathogenesis (“Tumor microenvironment pathway”) (Fig. 1E). Pathways enriched for downregulated genes were primarily related to cell cycle regulation (“Estrogen-mediated S phase entry”, “Cyclins and cell cycle regulation”, “Cell cycle G1/S checkpoint regulation”, “Cell cycle regulation by BTG family proteins”, “Cell cycle control of chromosomal replication”) (Fig. 1E). Intriguingly, viral pathogenesis-related pathways, specifically “Role of hypercytokinemia/hyperchemokinemia in the pathogenesis of Influenza” and “Role of PKR in IFN induction and antiviral response,” were among the most enriched pathways in our IPA analysis of upregulated genes. Together, these findings began to suggest that antiviral interferon expression, in addition to pro-inflammatory cytokines and chemokines, may play an important role in *R. equi* pathogenesis.

To more closely examine the macrophage innate immune response to *R. equi* infection, we first measured pro-inflammatory cytokine induction in macrophages infected with *R. equi*. At both 4- and 8 h, *R. equi*-infected macrophages had robust induction of the pro-inflammatory cytokines *Tnfa, Il1b* and *Il6*, consistent with activation of pattern recognition receptors such as TLR-2 (Fig. 2A). This was the case both for RAW 264.7 macrophages (Fig. 2A) as well as primary bone marrow-derived macrophages (BMDMs) (Fig. S1A). To track NF-κB activation, we measured phosphorylation of NF-κB in cell lysates by Western immunoblot analysis and observed robust NF-κB phosphorylation at 2-, 4- and 6h (Fig. 2B). Expression of these cytokines was at least partially dependent on MyD88; RAW 264.7 macrophages with *Myd88* knocked down by shRNA (*Myd88* KD, 65% efficiency) (Fig. S1B) had reduced levels of *Il1b, Il6* and *Tnfa* after *R. equi* infection compared to SCR (scramble shRNA) controls (Fig. 2C). These findings are consistent with a study by Darrah and colleagues, who reported TLR2- and MyD88-dependent production of pro-inflammatory (17).

**Figure 2:**
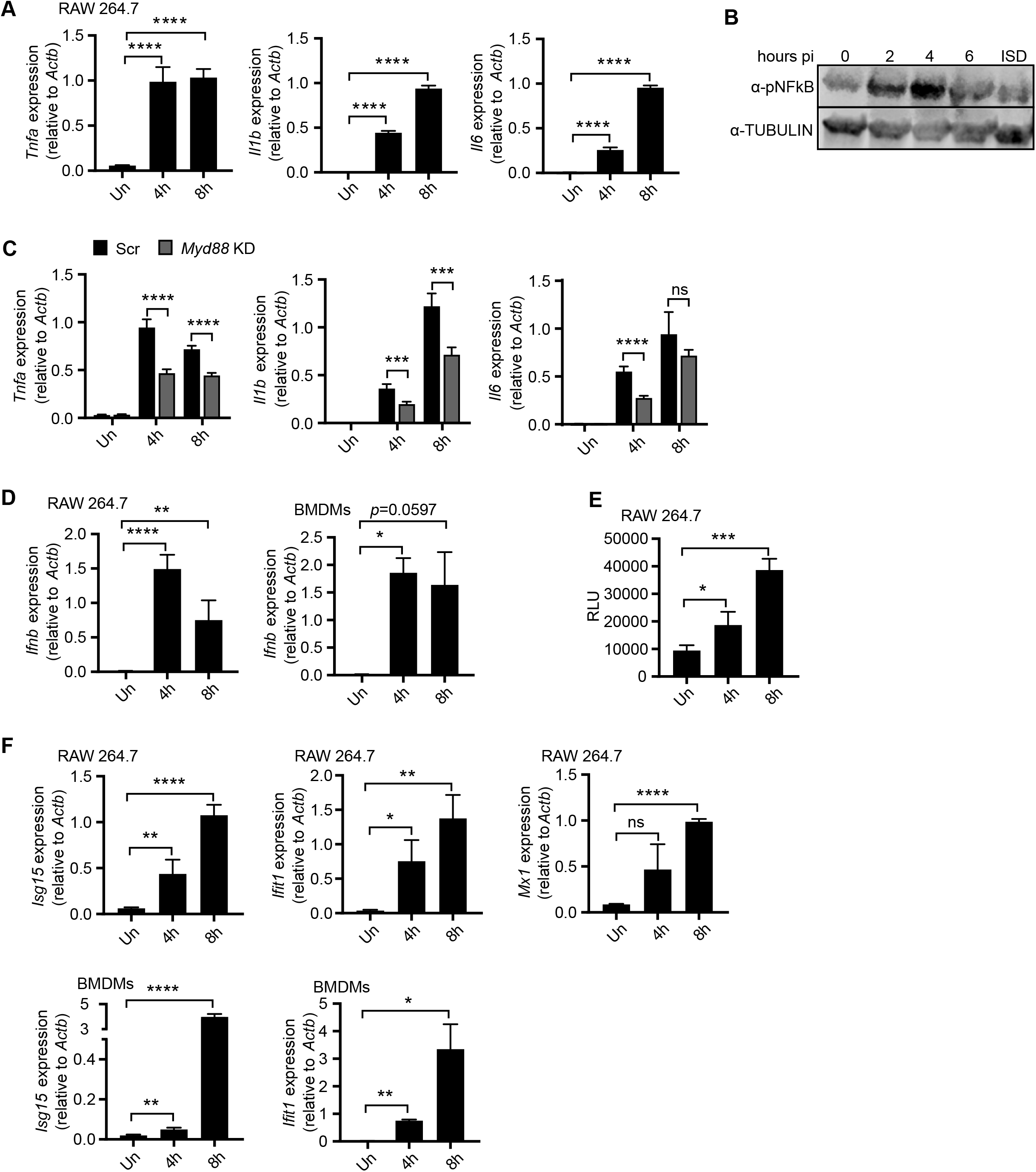
*R. equi* induces type I IFN expression during macrophage infection. (A) qRT-PCR of *Tnfa, Il1b* and *Il6* at indicated time post *R. equi* infection. (B) Western blot of pNFκB in *R-equi-infected* or ISD-transfected macrophages, with TUBULIN as a loading control. (C) qRT-PCR of *Tnfa, Il1b* or *Il6* in *R. equi-*infected Myd88 KD macrophages. (D) qRT-PCR of *Ifnb* in *R. equ*i-infected macrophages. (E) ISRE reporter cell assay with relative light units (RLU) measured as a readout for secreted type I IFN protein in *R. equi* infected macrophages. (F) As in (D) but of *Isg15, Ifit1* and *Mx1*. All qRT-PCRs represent 3 biological replicates ± SD, n=3. Statistical significance was determined using Students’ t-test. *p < 0.05, **p < 0.01, ***p < 0.001, ****p < 0.0001, ns = not significant.

Our transcriptomics analysis also revealed upregulation of several type I IFN response genes (Fig. 1C), and canonical pathway analysis revealed antiviral signaling as a highly enriched pathway in *R. equi*-infected cells (Fig. 1E). To further investigate the type I IFN response induced by *R. equi*, we first assessed the dynamics of *Ifnb* expression over a time-course of *R. equi* macrophage infection in murine BMDMs and in RAW 264.7 macrophages by RT-qPCR. We observed strong induction of *Ifnb* peaking at 4h post-infection in both cell types (Fig. 2D). To determine whether transcriptional upregulation of *Ifnb* led to elevated protein levels in *R. equi* infected cells, we used an IFN-stimulated response element (ISRE) luciferase reporter cell line as a readout for secreted IFN-α/β from *R. equi*-infected RAW 264.7 macrophages (Fig. 2E). Consistent with the transcriptional changes, we observed robust production of IFN-α/β protein in response to *R. equi* infection. Secreted IFN-β is recognized in an autocrine and paracrine manner through the IFN-β receptor (IFNAR1/2) and results in downstream expression of interferon stimulated genes (ISGs). Thus, we next measured expression of several ISGs, including *Isg15, Ifit1* and *Mx1*, over the same time course and observed significant induction of these genes 4h post-infection with peak induction at 8h in both cell types (Fig. 2F). Therefore, we concluded that *R. equi* infection elicits a robust type I IFN response in macrophages.

### The *R. equi* virulence factor VapA is not required for induction of pro-inflammatory cytokines or type I IFNs in macrophages

*R. equ?s* ability to survive and replicate within macrophages is largely dependent on a ~ 90 kb virulence plasmid and the virulence associated proteins (Vaps) it encodes. The best characterized, VapA, is required for virulence in foals and promotes intracellular survival (13, 15), as plasmid-cured strains of *R. equi* fail to replicate inside macrophages and do not cause inflammatory cell death (14). To investigate if VapA is required for inducing pro-inflammatory cytokines or type I IFNs in response to *R. equi* infection, we infected RAW 264.7 macrophages with isogenic strains of *R. equi* either with (33701+) or without (33701-) VapA at a MOI of 5 for 4- and 8h and measured gene expression. Macrophages infected with plasmid-cured *R. equi* (33701-) had reduced expression of *Tnfa* at 8h, and of *Il6* at both 4- and 8h after infection (Fig. 3A). Consistent with a previous study using *R. equi* strain ATCC 103 in mouse peritoneal macrophages (33), macrophages infected with *R. equi* 33701-had greater *Il1b* expression at 8h post-infection (Fig. 3F). However, *Ifnb* and *Isg15* expression in macrophages infected with *R. equi* 33701-was virtually identical to macrophages infected with virulent *R. equi* 33701+ (Fig. 3B). These findings indicate that while VapA does influence pro-inflammatory cytokine expression, it is not required for induction of type I IFNs during macrophage infections.

**Figure 3:**
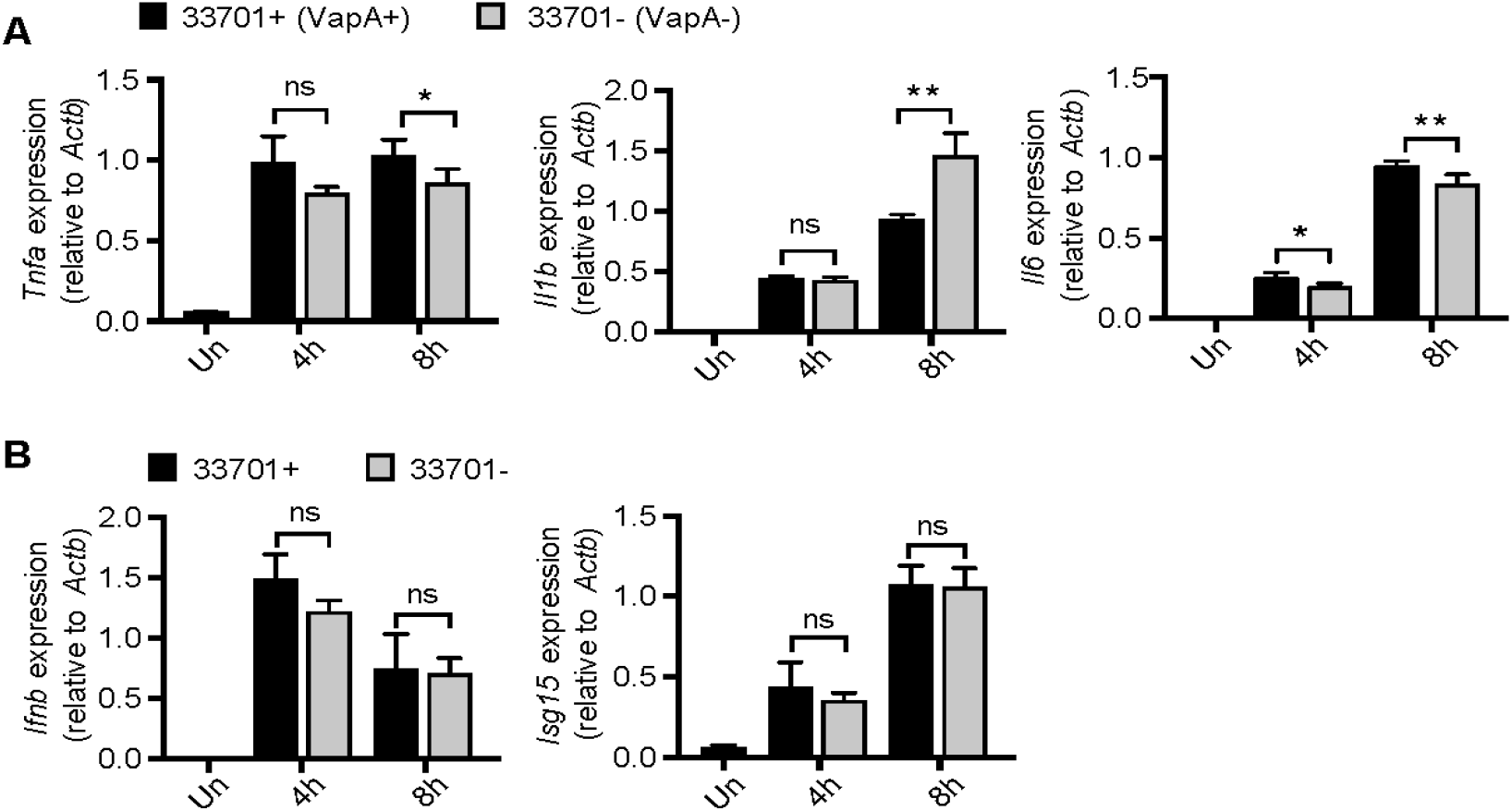
VapA is not required for induction of pro-inflammatory cytokines or type I IFN in murine macrophages. (A) qRT-PCR of *Tnfa, Il1b* and *Il6* in RAW 264.7 macrophages infected with *R. equi* with (33701+) and without (33701+) VapA. (B) As above but measuring *Ifnb* and *Isg15*. All qRT-PCRs represent 3 biological replicates ± SD, n=3. For all experiments in this study, statistical significance was determined using Students’ t-test. *p < 0.05, **p < 0.01, ***p < 0.001, ****p < 0.0001, n.s. = not significant.

### TBK1 is required for type I IFN induction in response to *R. equi* infection in primary murine macrophages

*Ifnb* is induced in a number of ways, including downstream of endosomal (e.g., TLR9 sensing of CpG DNA, TLR3 sensing of double-stranded RNA) or cytosolic (e.g., cGAS sensing of double-stranded DNA) sensing pathways, each of which triggers phosphorylation and activation of the IRF family of proteins, primarily IRF3 and IRF7. IRF7 is expressed at low levels in macrophages until induced downstream of type I IFN, IRF3 is expressed and activated of pathogen associated molecular patterns (34, 35). To determine if IRF3 is activated in response to *R. equi* infection, we infected RAW 264.7 cells (MOI of 50) and measured phosphorylated IRF3 (Ser396) by immunoblot. We detected IRF3 phosphorylation at 2-, 4- and 6h, peaking at 4h post-*R. equi* infection (Fig. 4A). We also examined STAT1 activation, which occurs downstream of the IFN a/b receptor IFNAR1/2, and found strong phosphorylation at 4, and 6h post-infection, with peak activity at 4h (Fig. 4A).

**Figure 4:**
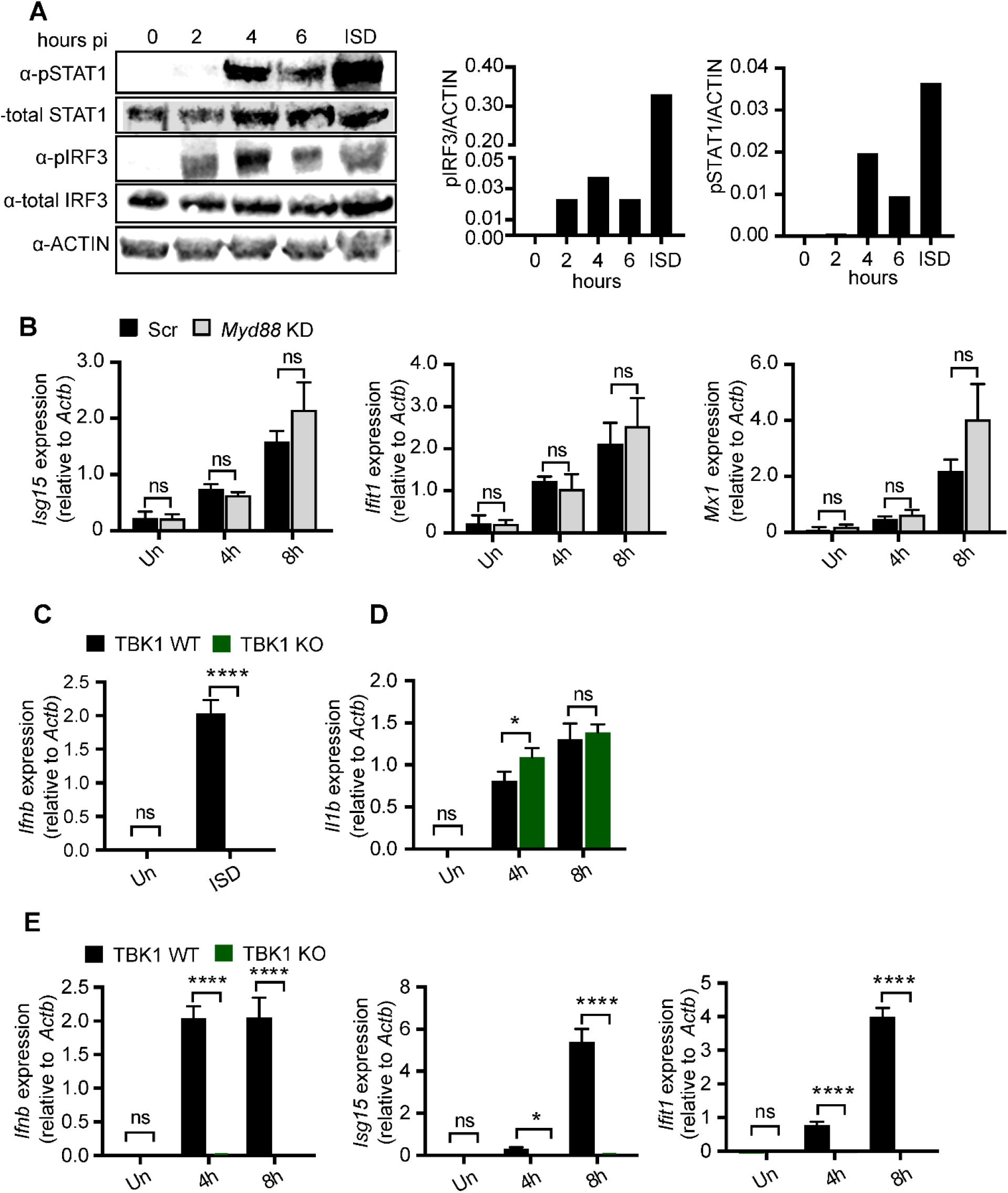
TBK1 is required for type I IFN induction in response to *R. equi* infection in primary murine macrophages. (A) Western blot of pSTAT1, total STAT1, pIRF3 and total IRF3 in RAW 264.7 macrophages infected or not (0) with *R. equi* for 2, 4 or 6 hours or transfected with ISD for 4 hours. ACTIN was used as a loading control. (B) qRT-PCR of *Isg15, Ifit1* and *Mx1* in MyD88 KD macrophages infected with *R. equi* for the indicated times. (C) qRT-PCR of *Ifnb* in control (TBK1^+/+^TNFR^-/-^) and KO (TBK1^-/-^TNFR^-/-^) BMDMs transfected with ISD for 4 hours. (D) qRT-PCR of *Il1b* in TBK1 BMDMs uninfected or infected with *R. equi* for the indicated times. (E) As in D but *Ifnb, Isg15* and *Ifit1*. All qRTPCRs represent 3 biological replicates ± SD, n=3. For all experiments in this study, statistical significance was determined using Students’ ttest. *p < 0.05, **p < 0.01, ***p < 0.001, ****p < 0.0001, n.s. = not significant.

Having observed activation of IRF3 in *R. equi*-infected macrophages (Fig. 4A), we hypothesized that the innate immune kinase TBK1 is required for type I IFN production. To test the contribution of TBK1 in type I IFN signaling during *R. equi* infection, we isolated BMDMs from mice lacking the kinase TBK1. Because *Tbk1* deletion in C57BL/6 mice causes a TNF receptor-dependent embryonic lethality, we used *Tbk1^-/-^/Tnf^-/-^* double KO (TBK1 KO) mice to assess the contribution of TBK1 and compared them to *Tbk1^+/+^/Tnf^-/-^* controls (28, 36). As expected, *Tbk1^-/-^* macrophages failed to induce *Ifnb* in response to transfection with the double-stranded DNA agonist ISD (37) (Fig. 4C). *R. equi*-infected *Tbk1^-/-^*BMDMs had no difference in expression of the pro-inflammatory cytokine *Il1b* compared to controls (Fig. 4D). However, *R. equi*-infected *Tbk1^-/-^* macrophages had almost no induction of *Ifnb, Isg15* and *Ifit1* at 4- and 8h after infection (Fig. 4E), indicating that TBK1 is required for induction of a type I IFN response following infection with *R. equi*.

Because autocrine sensing of IFN-β by IFNAR1/2 elicits ISG expression via STATs, we predicted that loss of IFNAR would abrogate the type I IFN response in *R. equi* infected macrophages. Compared to WT controls, the *Ifnar1^-/-^* macrophages had no differences in pro-inflammatory cytokine levels after *R. equi* infection (Fig. S2A). However, *R. equi*-infected *Ifnar1^-/-^* macrophages displayed a modest reduction in *Ifnb* levels (Fig. S2B), and ISG expression (*Isg15*) was completely abrogated (Fig. S2B). As a positive control, when we transfected *Ifnar1^-/-^* macrophages with ISD (which stimulates *Ifnb* production), there was no ISG induction (Fig. S2C).

We next investigated specific adapter proteins in type I IFN induction during *R. equi* infection. Type I IFN production downstream of TLRs is mediated via MyD88/TRIF, while cytosolic DNA and RNA sensing signals through cGAS/STING and MAVS/RIG-I, respectively. We first measured ISG expression (i.e., *Isg15*, *Ifit1* and *Mx1*) in *Myd88*- and *Trif*-knockdown macrophages (*Trif* KD, 70% efficiency) (Fig. 4B, S2D) and detected no major differences in ISG expression with the knock down of either TLR adapters, with the exception of a slight reduction in *Isg15* expression in *Trif* knockdown RAW 264.7 cells (Fig. S2E). These results suggest that neither TLR9 nor TLR3 are required for type I IFN signaling during *R. equi* infection. Collectively, these results demonstrate that that *R. equi* induces type I IFN through TBK1 and IFNAR, but that this occurs via a sensor other than TLR3 or 9.

### Cytosolic DNA sensing via cGAS/STING/TBK1 is required to induce type I IFN during in macrophages infected with *R. equi*

Having ruled out type I IFN induction by several adapter proteins downstream of TLR sensors, we next asked whether *R. equi* infection directly stimulates cytosolic nucleic acid sensing. STING is an adaptor protein in the cytosolic DNA sensing axis that is activated by host cyclic dinucleotides like cGAMP produced by activated cGAS or by bacterial cyclic dinucleotides like c-di-AMP or c-di-GMP (38–40). To investigate whether STING is required for *R. equi* induction of type I IFNs, we infected CRISPR-Cas9-generated STING knock out (KO) RAW 264.7 macrophages with *R. equi* and measured cytokine expression by RT-qPCR. In response to *R. equi* infection, while STING KO macrophages induced *Tnfa* to comparable levels as control macrophages (expressing a GFP-targeting guide RNA), they failed to induce *Ifnb*, *Isg15*, or *Ifit1* at 4- or 8h post-infection (Fig. 5A). As a positive control we transfected STING KO RAW 264.7 macrophages with ISD (to stimulate STING via cGAS) and found that these cells fail to induce *Ifnb* (Fig. S3A) (41). Using ISRE reporter cells, we also observed a reduction in the protein levels of IFN-α/β in the supernatants of STING KO macrophages compared to controls (Fig. S2B). These data indicate that STING is absolutely required for type I IFN expression in response to *R. equi* infection in macrophages.

**Figure 5.**
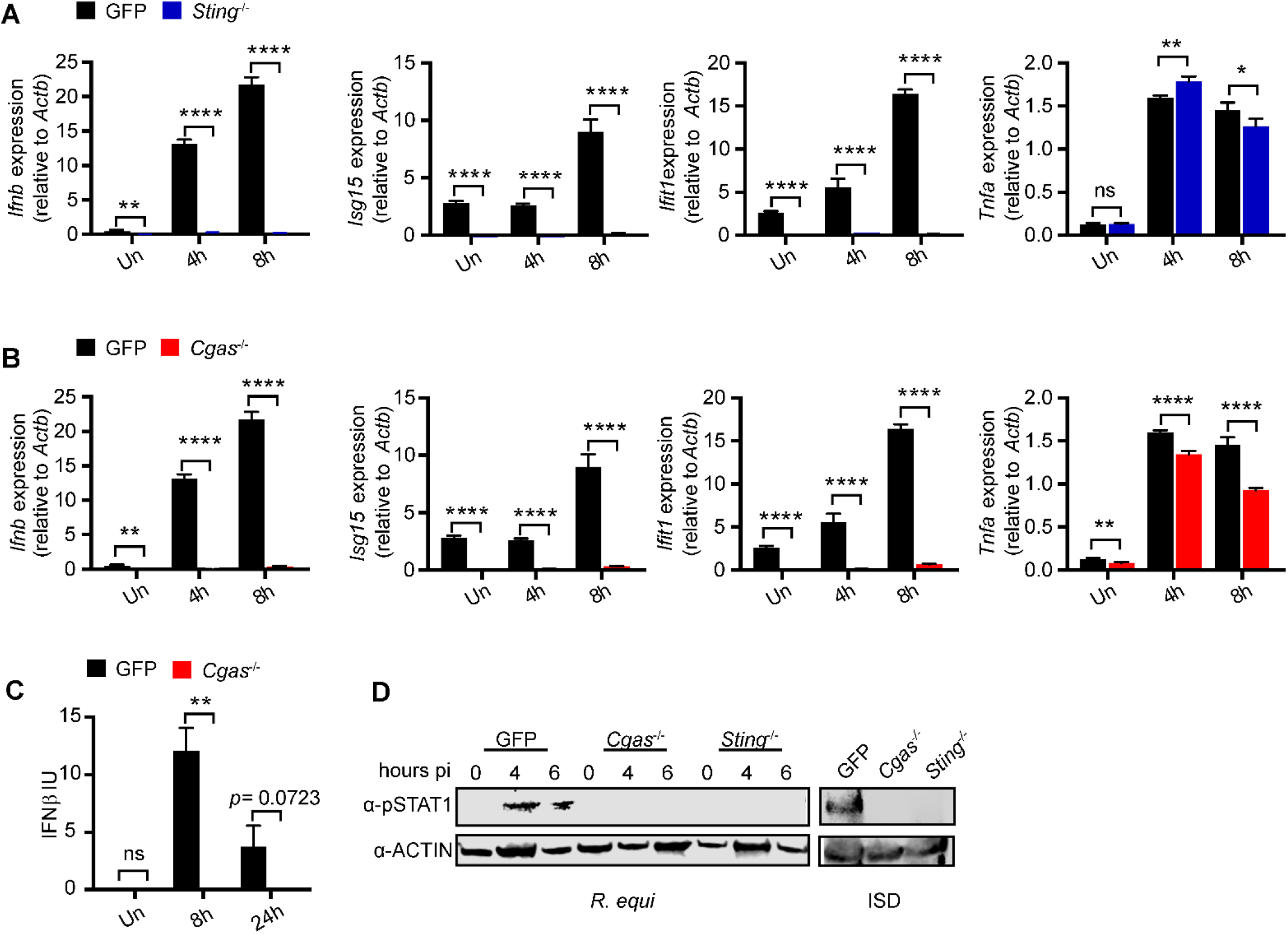
The cytosolic DNA sensing axis of cGAS/STING/TBK1 is required to induce type I IFN during *R. equi* infection of macrophages. (A) qRT-PCR of *Ifnb, Isg15, Ifit1* and *Tnfa* in GFP control or STING KO RAW 264.7 macrophages infected with *R. equi* for the indicated times. (B) As in A but in GFP control and cGAS KO RAW 264.7 macrophages. (C) IFN-β protein ELISA of RAW 264.7 macrophages infected with *R. equi* for the indicated times. (D) Western blot of pSTAT 1 in GFP, cGAS KO and STING KO RAW 264.7 cells infected *R. equi* for the indicated times or transfected with ISD for 4 hours. ACTIN was used as a loading control. All qRT-PCRs represent 3 biological replicates ± SD, n=3. For all experiments in this study, statistical significance was determined using Students’ t-test. *p < 0.05, **p < 0.01, ***p < 0.001, ****p < 0.0001, n.s. = not significant.

Some intracellular bacterial pathogens such as *Listeria monocytogenes* activate cytosolic sensing by producing cyclic dinucleotides (c-di-AMP) that bind to and activate the adaptor protein STING, which in turn activates TBK1 (42). To determine if *R. equi* might activate the cytosolic DNA sensing pathway in a similar way, we searched the Kegg database and found that *R. equi* encodes a cyclic dinucleotide, diadenylate cyclase that could produce c-di-AMP (43). To test if *R. equi* was stimulating STING directly via production and secretion of c-di-AMP or via activation of cGAS, which produces the cyclic dinucleotide cGAMP in response to binding cytosolic double-stranded DNA (44, 45), we generated cGAS KO RAW 264.7 cells using CRISPR-Cas9. As with loss of STING, cGAS KO macrophages fail to induce *Ifnb* in response to ISD transfection (Fig. S3C). Upon infection with *R. equi*, cGAS KO macrophages had reduced levels of *Ifnb, Isg15* and *Ifit1* transcripts (Fig. 5B), and a significant but less dramatic reduction in *Tnfa* (Fig. 5B). At the protein level, cGAS KO macrophages had defective IFN-α/β production as measured by ISRE reporter cells, but notably, expression was not completely ablated (Fig. S3D). However, when we measured IFN-β protein levels at 0-, 8- and 24h postinfection by ELISA, we observed a complete loss of IFN-β in supernatants from both cGAS and STING KO macrophages (Fig. 5C). This loss of type I IFN production was not due to impaired bacterial engulfment or replication as bacterial burden was similar in control, STING KO and cGAS KO macrophages (Fig. S3E).

Because there was a slight inconsistency in IFN-β levels measured by ISRE reporter cells (which measure both IFN-α and −β) and by an IFN-β-specific ELISA (Fig. 5C, S3D), we also measured STAT1 activation in *R. equi*infected control, STING KO and cGAS KO macrophages by immunoblot, probing for phosphorylated (activated) STAT1 (Tyr701). We found that in control macrophages STAT1 was robustly phosphorylated at 4- and 6h post-infection, phosphorylated STAT1 was undetectable in STING KO or cGAS KO macrophages at all time points (Fig. 5D). These results show that *R. equi* infection results in activation of the cGAS/STING/TBK1 axis and further suggests that release of double stranded DNA into the cytosol is responsible for activation of the cytosolic DNA sensing pathway leading to production of the type I IFN.

### Galectin-3, −8, and −9 are recruited to *R. equi* containing vacuoles

Because *R. equi* is confined by a phagosomal membrane until 24h post-infection in macrophages (46), it was puzzling to observe activation of the cytosolic DNA sensor cGAS early during infection. Previous studies discovered that Mtb, another “vacuolar” pathogen, permeabilizes its phagosomal membrane using its ESX-1 secretion system (31). Therefore, we hypothesized that *R. equi* may also permeabilize its phagosomal membrane to allow for communication with the host cytosol, resulting in detection by cytosolic DNA sensors. We recently reported that the cytosolic glycan-binding proteins galectin-3, −8, and −9 access the lumen of damaged Mtb-containing vacuoles following ESX-1-mediated permeabilization (47). To investigate whether *R. equi* phagosomal membranes are sufficiently damaged during infection to recruit these galectins, we infected RAW 246.7 cells stably expressing 3xFLAG-tagged galectins (47). Specifically, we looked at galectins-3, −8, −9 because they have been shown to colocalize with Mtb, *L. monocytogenes*, *Salmonella enterica* serovar Typhimurium, and *Shigella flexneri* (47, 48); galectin-1 was used as a negative control. Each of these galectin cell lines were infected with virulent GFP-expressing *R. equi* (MOI 5) and over a time course of 4-, 8-, and 16h, cells were fixed and imaged by immunofluorescence microscopy (49). Galectin-3, −8, and −9 but not galectin-1 were all recruited to *R. equi* to varying degrees. Recruitment of galectins-8 and −9 peaked at 8h post-infection, with galectins-8 and −9 recruited to ~10% and ~6% of *R. equi*, respectively (Fig. 6A, B). Galectin-3 recruitment peaked at 4h post-infection, with recruitment to ~12% of *R. equi* at 4h and declining to ~5% by 16h. Curiously, by 4h post infection, galectin-3, −8, and −9 formed puncta in the cytosol of *R. equi* infected cells (Fig. 6C), but the nature of these puncta is unclear. Galectin-1 formed rare puncta but did not associate with *R. equi* (Fig. 6C). A galectin-positive population of *R. equi* suggests that phagosomal membrane damage and access to the cytosol occurs as early as 4h following infection. Interestingly, at 16h, many of the galectin-positive *R. equi* had markedly reduced GFP expression compared to galectin-negative bacteria, suggesting they had potentially been killed or lysed.

**Figure 6.**
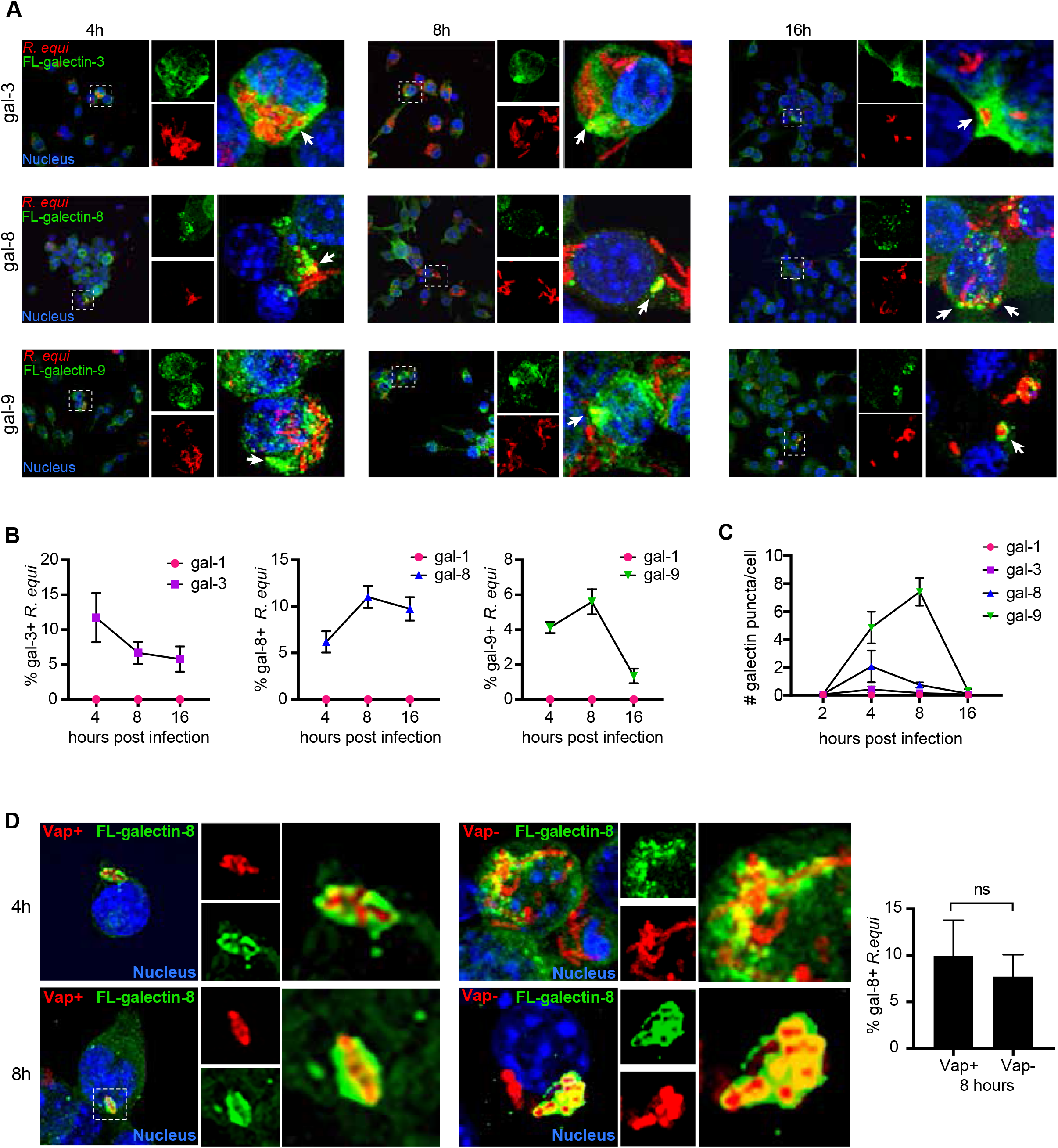
Galectin-8 is recruited to *R. equi*. (A) Immunofluorescence (IF) of RAW 264.7 cells stably expressing 3XFLAG (FL)-tagged galectin-3, −8, or −9 infected with GFP expressing *R. equi* 103+ (MOI-5) for the indicated times. (B) Quantification of galectin-1, −3, −8, or −9 recruitment to R. equi at the indicated times. (C) Quantification of galectin-1, −3, −8, or −9 puncta in *R. equi*-infected RAW 264.7 cells at the indicated times. (D) IF of RAW 264.7 cells stably expressing 3X FL-tagged galectin-8 infected with GFP expressing *R. equi* 103+ or 103- (MOI-5) for the indicated time and quantification of galectin-8 positive *R. equi* in at 8 hours expressed as percent of total *R. equi*. IF images are representative of at 3 independent experiments. Quantification is the percent of positive bacteria with at least 100 bacteria quantified per coverslip. Error bars are ±SEM. Statistical significance was determined using Students’ t-test. *p < 0.05, n.s. = not significant.

Given that the *R. equi* virulence factor VapA has been shown to permeabilize lysosomal membranes to modulate lysosome pH (13) and required for survival and replication within macrophages (14), we hypothesized that VapA might be required for permeabilizing the phagosomal membrane and permitting galectin recruitment. To test this, we infected 3xFLAG tagged galectin-8 RAW246.7 cells with GFP-expressing, plasmid-cured *R. equi* 103-, which does not express VapA (50). The plasmid cured strain of *R. equi* had similar recruitment of galectin-8 as VapA expressing *R. equi* (Fig. 6D), indicating that the *R. equi* phagosomal membrane permeabilization does not require VapA and occurs via an alternative mechanism. Taken together, these data indicate that the *R. equi*-containing vacuole is permeable and accessible to the macrophage cytosol, which could enable detection of bacterial-derived ligands by cGAS and other cytosolic sensors.

### *R. equi* induces both a pro-inflammatory and type I IFN expression program *in vivo*

Having shown that *R. equi* induces type I IFNs in macrophages, we next sought to determine how the type I IFN response contributes to *R. equi* pathogenesis *in vivo*. Because we can exert a greater degree of control over the murine environment, we began by infecting 8-week-old C57BL/6 mice with *R. equi* by intraperitoneal (IP) injection. At 5d post-infection, we harvested lungs, mesenteric lymph nodes, spleens and peritoneal cells (Fig. 7A). Age-matched mice injected with an equal volume of PBS (Un) served as negative controls. Bacteria were recovered from the spleen, mesenteric lymph nodes and lungs, which indicates the bacteria disseminated from the point of initial infection (Fig. 7B). We measured spleen weights as a readout for inflammatory responses and observed a ~1.5-fold increase in splenic weight in infected mice compared to those injected with PBS (Fig. 7C). We measured pro-inflammatory cytokines and type I IFNs in the spleen and peritoneal cells isolated from *R. equi*-infected mice 5d post-infection by qRT-PCR, and consistent with our observations in macrophages, spleens and peritoneal cells from mice infected with *R. equi* had significantly elevated levels of both pro-inflammatory cytokines and ISGs compared to uninfected controls (Fig. 7D)

**Figure 7:**
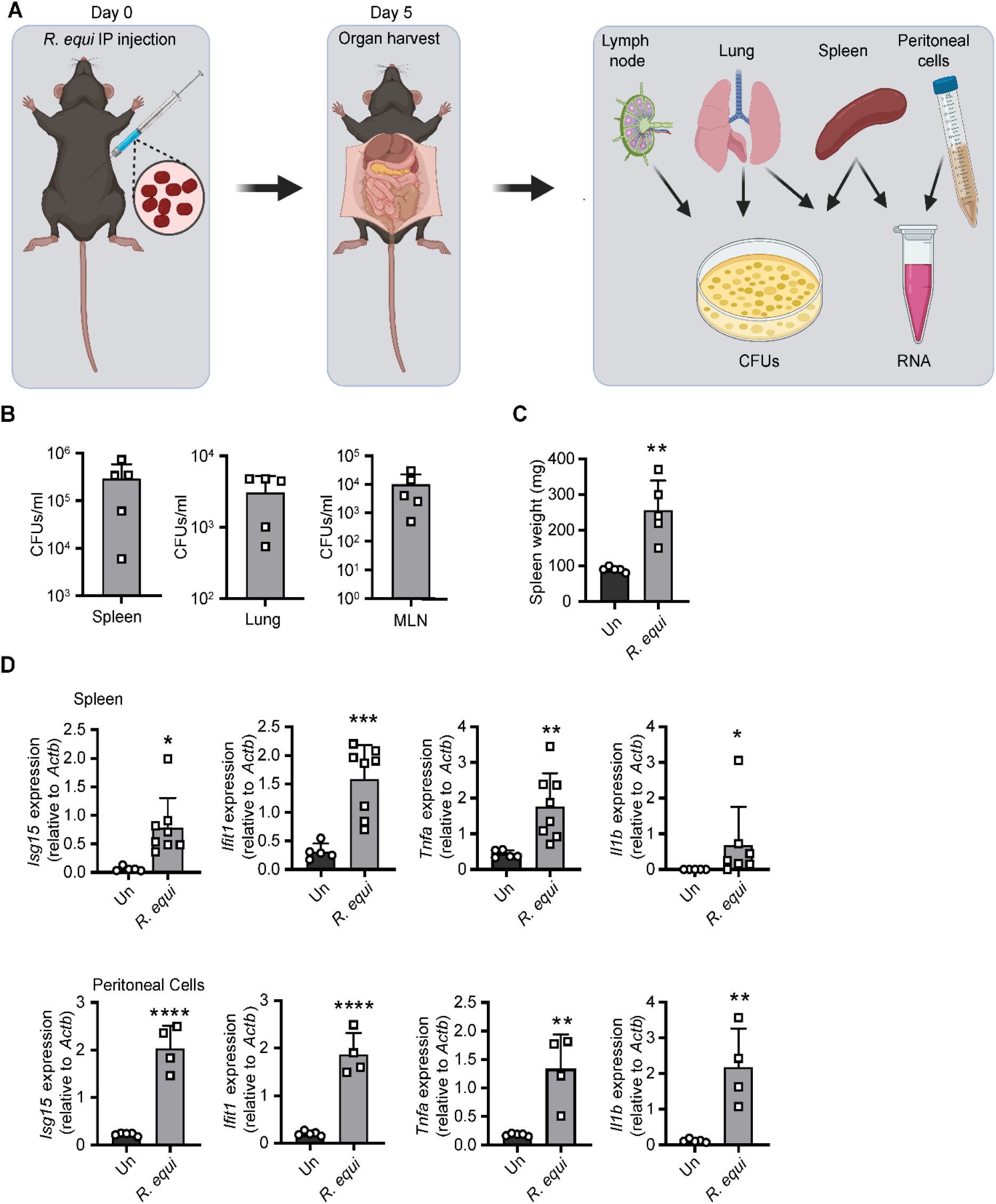
*R. equi* induces both a pro-inflammatory and type I IFN expression program in a mouse model of infection. Figure 7 R. equi induces both a pro-inflammatory and type I IFN expression program in a mouse model of infection. (A) Mouse R. equi infection schematic. Mice were infected with R. equi by IP injection on Day 0. On day 5, organs were harvested for CFU and qRT-PCR analysis. (B) CFUs/gram of the spleen, lung and mesenteric lymph nodes of mice 5d post-infection. Each square represents an individual mouse. (C) Spleen weight in mg of mice 5d post R. equi infection. (D) qRT-PCR of Isg15, Ifit1, Tnfa and Il1b in the spleen or peritoneal cells of mice uninfected or infected with R. equi. Mouse qRT-PCRs represent 5 biological replicates ± SD. For all experiments in this study, statistical significance was determined using Mann-Whitney test. *p < 0.05, **p < 0.01, ***p < 0.001, ****p < 0.0001, ns = not significant.

Because mice are relatively resistant to *R. equi* infection, and since horses are the primary natural host and most physiologically relevant model, we next turned to an equine model infection. We collected pre-infection, baseline blood samples from 28-day-old horses and assessed their lungs for pulmonary lesions by thoracic ultrasonography. We next infected foals intrabronchially via endoscopy (51–53) with 1 x 10^6^ virulent *R. equi*, and monitored them for 3 weeks; age-matched uninfected foals served as negative controls. At 21d post-infection, we again collected blood and evaluated lungs by thoracic ultrasonography to monitor disease progression or resolution. (Fig. 8A). To investigate if *R. equi* induces a type I IFN program in horses, we isolated monocytes from peripheral blood samples taken at 0- and 21d post-infection and measured ISGs by RT-pPCR. To rule out age-related changes in ISG levels, we first compared uninfected and *R. equi* infected foals at 21d post-challenge, normalizing individual post-challenge values to baseline levels. Uninfected foals had no significant induction of ISGs between baseline and 3 weeks post-challenge samples (Fig. 8B). However, we observed a significant induction of *ISG15* in infected foals, with infected foals falling into subgroups with modest induction or high induction. Notably, the sole foal that became clinically ill following experimental challenge fell within the high induction subgroup (denoted by yellow square in Fig. 8B). Since individual foals varied in their response to infection, and having shown that pre- and post-challenge cytokine levels were similar in uninfected foals, we next focused on infected foals and compared baseline *ISG15* levels with post-infection levels. We observed an upward trend in *ISG15* induction compared to pre-infection samples, but it did not reach statistical significance (Fig. 8C). We also observed a robust induction of pro-inflammatory cytokines (*IL1B, IL6*) in *R. equi*-infected foals (Fig. 8D). Given that *R. equi* is ubiquitous in the environment and likely shed by mares, and given that 2 foals exhibited evidence of pulmonary lesions prior to experimental infection (denoted by red circles in Fig. 8C-D), it is possible that foals were exposed to environmental *R. equi*, which may have contributed to innate immune activation during the 0d blood collection. Nonetheless, these results indicate that *in vivo R. equi* infection results in type I IFN activation in both mouse and equine models of infection.

**Figure 8:**
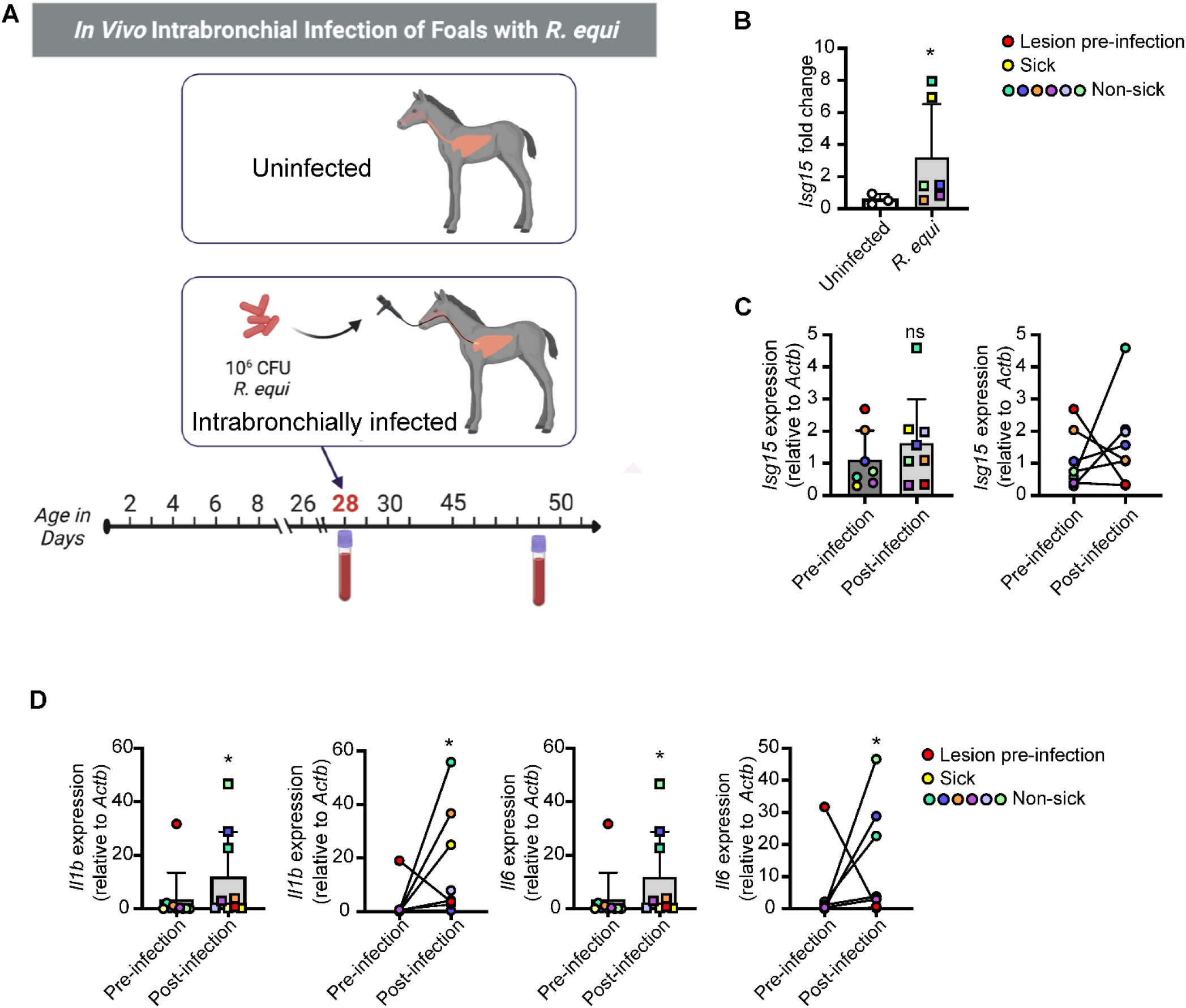
*R. equi* induces type I IFN response in equine monocytes (A) Horse *R. equi* infection schematic. Baseline blood samples were taken at 28 days of age. 28-day-old foals were then infected with *R. equi* and monitored for 3 weeks. 21 days post infection (at 49 days of age), blood was again harvested and a thoracic ultrasound was performed. Peripheral blood derived monocytes were processed for RNA and used for qRT-PCR analysis. (B) qRT-PCR of Isg15 in sham infected and *R. equi* infected foals at 3 weeks post infection normalized to baseline Isg15 levels. Each dot represents an individual foal. (C) As in B, but comparing baseline Isg15 levels to 3 weeks post infection in infected foals. Red dot indicates foal with lesions prior to experimental infection and yellow dot indicates foal that became clinically ill after infection. (D) As in C, but for Il1b and Il6. Horse qRT-PCRs represent >3 biological replicates ± SD, uninfected n=3, infected n=6. For all experiments in this study, statistical significance was determined using Mann Whitney test. *p < 0.05, **p < 0.01, ***p < 0.001, ****p < 0.0001, n.s. = not significant

## DISCUSSION

The innate immune response is critical as a first line of defense in limiting pathogen replication, but also shapes the adaptive immune response. Because innate immune responses dictate adaptive immune outcomes, defining the innate immune milieu generated in response to bacterial pathogens like *R. equi* is a crucial step to understanding how it causes disease. Here we show that intracellular *R. equi* triggers the cytosolic DNA sensing pathway in a cGAS- and STING-dependent manner. Activation of the cytosolic DNA sensing pathway following *R. equi* infection led to phosphorylation of IRF3 and initiation of a type I IFN response within 4h of infection, while intracellular bacteria are still replicating within the *R. equi*-containing vacuole (16, 54). Our finding that this type I IFN expression profile is dependent upon the cGAS/STING/TBK1 axis supports a model whereby *R. equi* persists in a modified vacuole capable of interacting with the macrophage cytosol and engaging DNA sensors and other innate immune proteins.

Our observation that vacuolar *R. equi* induces type I IFN expression via activation of the DNA sensing pathway parallels that of Mtb in macrophages. Mtb uses its type VII secretion system, ESX-1, to permeabilize its vacuolar membrane, permitting liberation of extracellular mycobacterial DNA into the macrophage cytoplasm where it triggers cGAS (28). Genetic evidence indicates that like Mtb, *R. equi* also encodes a type VII secretion system, although the function of this system in *R. equi* remains to be determined (43, 55). Further studies focused on investigating the function of *R. equi*’s ESX secretion systems will be important for uncovering the mechanism of cytosolic DNA sensing during macrophage infection. Mtb ESX-1 effectors ESAT-6 and CFP-10, together with the mycobacterial lipid PDIM, are thought to form pores in the phagosome after their secretion (56, 57); the presence or function of similar effectors in *R. equi* might suggest a similar mechanism, but *R. equi* could have unrelated ways of accessing the cytosol to communicate with and manipulate the host. Using readouts and phenotypes we identified here, future screens may serve to elucidate these mechanisms and uncover novel *R. equi* effectors and effector functions.

The recruitment of the glycan binding proteins galectin-3, −8, and −9 to a population of *R. equi* indicates that the *R. equi*-containing vacuole is damaged to the extent that luminal glycans are exposed and recognized by cytosolic danger sensors as early as 4h post infection. This is consistent with Mtb recruitment of these galectins, which occurs as early as 3h post-infection (47), while *Salmonella enterica* serovar Typhimurium recruits galectin-3, −8, and − 9 to broken vacuoles as early as 1h post-infection (48). Interestingly, we observed an accumulation of galectin puncta that are unassociated with bacteria. It is possible that virulence factors responsible for permeabilizing *R. equi*’s phagosome are secreted and act in trans to also damage endosomal compartments. Further studies will have to be conducted to understand the nature and fate of the damaged phagosomes and endosomes.

Our finding that cGAS is required for induction of type I IFN signaling during *R. equi* infection indicates that DNA is the major contributor for this activation in macrophages; however, the source of this DNA remains to be determined. Given the data supporting damaged *R. equi* phagosomes, it is likely that bacterial DNA is released into the cytosol, however the possibility of mitochondrial perturbation and cytosolic release of mitochondrial DNA has not been ruled out. Indeed, recent evidence indicates that Mtb infection induces release of host mitochondrial DNA into the cytosol, contributing to the type I IFN response during infection (58, 59). Additionally, the cyclic dinucleotide produced by *L. monocytogenes* and Mtb, (cyclic diadenosine monophosphate (c-di-AMP)), bypasses cGAS and its second messenger cGAMP to directly activate STING (60). While some bacteria can directly activate STING, through secretion of c-di-nucleotides, and while *R. equi* does seem to encode deadenylate cyclases, our data suggest that this is not the mechanism of type I IFN induction under the conditions or time points we examined. Because cGAS KOs do not induce type I IFNs, it indicates that without host derived cGAMP, no bacterial CDNs are present to activate STING and induce type I IFNs. It is now clear that intracellular bacterial pathogens have evolved to elicit type I IFN through activating the cytosolic DNA sensing pathway including cGAS (Mtb (28, 61),*Chlamydia* spp. (44), *F. novicida* (27)) and/or STING (*L. monocytogenes* (62), Mtb (63),and *Chlamydia trachomatis* (64)) implicating a selective advantage for at least some bacteria to elicit this response, likely by ISG activity such as blocking IL-1 activity via ILRα (65), or IL-10 mediated limitation of IFN-γ responses (24).

A key virulence determinant of *R. equi* is the conjugative virulence plasmid which hosts a 21-kb pathogenicity island encoding Vaps. The best characterized of the Vaps is VapA, which is required but not solely sufficient for intracellular growth within macrophages (14, 66). VapA promotes *R. equi* survival in macrophages by inhibiting phagosomal maturation (50), inducing lysosome biogenesis (15), and contributing to lysosome membrane permeabilization (13). Interestingly, disruption of the phagosomal membrane, activation of cytosolic DNA sensing, and production of type I IFNs was not dependent on expression of Vap A. This discrepancy may be explained by the transient nature or small size of VapA induced membrane leaks, with VapA-induced membrane lesions ranging from 0.37 nm at pH 6.5 to 1.05 nm at pH 4.5, which are large enough for the passage of ions (13). Additional studies investigating additional bacterial mutants may help elucidate addition bacterial factors underlying in the hostpathogen interactions of *R. equi* and macrophages.

It will also be interesting to interrogate downstream outcomes of *R. equi* infection and pathogenesis that result from the activation of cytosolic sensing, especially bacterial survival and replication in macrophages. One potential outcome of TBK1 activation during *R. equi* infection is autophagic targeting (28, 30, 67). Autophagy, and selective autophagy in particular, functions as an antimicrobial mechanism that promotes lysosomal degradation of intracellular bacterial pathogens (68). Future studies will be needed to investigate if *R. equi* also gets targeted to this pathway, but its recruitment of galectins and activation of TBK1 suggests that a population of *R. equi* could potentially be targeted and destroyed by this mechanism. VapA, while seemingly unimportant for type I IFN signaling, is important for evading lysosomal degradation (13) so it may provide a mechanism to counteract the potential drawbacks for bacteria activating this anti-bacterial pathway.

Another outstanding question is how induction of type I IFN signaling influences *R. equi* pathogenesis. It may be interesting to determine the transcriptional signature in foals with severe disease to test the hypothesis that foals that succumb to infection have a robust, or even hyperinduced, type I IFN signature. While we observed induction of type I IFN in response to *R. equi* infection during our equine experiments, we were unable to draw clear conclusions regarding connections between type I IFN levels and disease outcome largely due to the small study size. All animals in the equine study ultimately cleared infection without treatment, and only one foal became clinically ill following experimental challenge; however this animal did have highly induced ISGs. Future studies in both mouse and horse models specifically centered on the balance between type I and type II IFN will be key in understanding the clinical implications of a type I IFN-skewed response, and whether such a signature correlates with reduced IFN-g, impaired control of bacterial replication, or detrimental disease outcomes. Identifying factors associated with negative disease outcomes will help determine ways to expedite diagnoses, promote positive disease outcomes, and reduce patient mortality.

## METHODS

### Animals

All experiments for this study were reviewed and approved by the Texas A&M University Institutional Animal Care and Use Committee. Mice were kept on a 12h light/dark cycle and provided food and water *ad libitum*. Mice were group housed (maximum 5 per cage) by sex on ventilated racks in temperature-controlled rooms. TBK1 KO mice (*Tnfr1^-/-^* and *Tbk1^-/-^/Tnfr1^-/-^*) (69) were generously provided by the Akira lab.

### Cell culture

Bone marrow-derived macrophages (BMDMs) were differentiated from bone marrow cells isolated by washing mouse femurs with 10 ml DMEM. Harvested cells were centrifuged for 5 min at 1000 rpm and resuspended in BMDM media ((DMEM, 20% FBS, 1 mM Sodium pyruvate, 10% MCSF conditioned media). Cells were counted and plated at 5 x 10^6^ in 15 cm non-TC treated dishes in 30 ml complete media and fed with an additional 15 ml of media on Day 3. On Day 7, cells were harvested with 1x PBS-EDTA. RAW 264.7 cells were purchased from ATCC and the cell line was minimally passaged in our laboratory to maintain genomic integrity, and new cell lines generated from these low passage stocks. Cell lines were passaged no more than 10 times and tested negative for mycoplasma contamination. Cells were cultured in complete media (DMEM, 10% FBS, 2% HEPES buffer).

### Monocyte isolation

Thirty ml of heparinized blood was incubated at room temperature for 30 min. The plasma layer containing white blood cells was removed, diluted with the same volume of PBS, and layered over Ficoll-Paque™ Plus (GE Healthcare, Uppsala, Sweden) for density gradient separation of peripheral blood mononuclear cells (PBMCs). PBMCs were washed 3 times with PBS, counted in an automated cell counter (Cellometer Auto T4, NexelomBioscience, Lawrence, MA), and suspended at a concentration of 3 × 10^6^cells/mL in RPMI-1640 (BioWhittaker^®^, Lonza, Walkersville, MD, USA) containing 15% heat-inactivated horse serum, 1% Glutamax™ (Life Technology Corporation, Grand Island, NY, USA), 1% NEAA mixture (BioWhittaker^®^, Lonza, Walkersville, MD, USA), penicillin G (100 U/mL), and streptomycin (80 μg/mL). PBMCs were incubated at 37 °C and 5% CO_2_ for 3h in a T75 flask, and then non-adherent cells were removed with warm PBS. Adherent cells were detached from the flask with Accutase^®^ (Innovative Cell Technology) for 10 min at room temperature and PBS. Cells were pelleted, washed with PBS to remove Accutase^®^, and the adherent cells (monocytes) counted using the automated cell counter, pelleted again, and stored in Trizol^®^ (Invitrogen) at −80 °C until RNA extraction.

### Stimulations

RAW 264.7 cells and murine BMDMs were plated in 12 well dishes at 5×10^5^ cells/well and allowed to adhere overnight. Cells were stimulated with 100 ng/ml LPS or transfected with 1 μg/ml ISD for 4h.

### Bacterial strains

Unless otherwise noted, virulent *Rhodococcus equi* 33701+ (VapA +; ATCC reference strain; Rockville, MD) was used as the wild type strain. The plasmid cured (33701-) strain was previously described (70). *R. equi* ATCC 103 expressing GFP (49, 50) were generously provided by Dr. Mary Hondalus.

### Macrophage infections

One colony forming unit (CFU) of *R. equi* was inoculated into 5 ml of brain-heart infusion (BHI) broth and shaken overnight at 37° C, then 0.5 ml of the culture was subcultured into 5 ml of fresh BHI overnight, shaken at 37°C. The bacterial suspension was centrifuged at 1000 rpm for 10 minutes at 25°C. The suspension was discarded and the pellets washed twice with 1 ml of phosphate buffered saline (PBS). The supernatant was discarded, the bacteria re-suspended in sterile PBS, and the concentration of bacteria determined spectrophotometrically at an optical density of 600 nm (OD 600) where OD 1.0 represents approximately 2×10^8^ CFU/ml. Concentrations were verified by plating the inoculum and counting CFUs. Bacterial suspensions were diluted to the desired concentration. Strains were confirmed to be virulent (VapA positive) by PCR before infection. Macrophages were plated in 12-well dishes at 5×10^5^ cells/well and allowed to adhere overnight. Macrophages were infected with virulent *R. equi* at a MOI of 5 for gene expression studies, MOI of 1 for CFU experiments or MOI of 50 for Western immunoblot analysis. Following infection, macrophages were centrifuged for 10 minutes at 1000 rpm, then incubated for 30 minutes at 37°C. Noninfected macrophage monolayers were cultured under the same conditions. Media containing *R. equi* was removed and each well washed twice with PBS, then replaced with complete media and cultured at 37°C until the appropriate time point. At each time point, supernatants were removed and monolayers were washed with PBS.

### Experimental infection of foals

For transendoscopic infection (51–53), foals were sedated using intravenous (IV) injection of xylazine hydrochloride (0.5 mg/kg; Vedco, St. Joseph, MO) and IV butorphanol tartrate (0.02 mg/kg; Zoetis, Florham Park, New Jersey). An aseptically-prepared video-endoscope with outer diameter of 9 mm was inserted via the nares into the trachea and passed to the bifurcation of the main-stem bronchus. A 40-mL suspension of virulent EIDL 5–331 *R. equi* containing approximately 1 x 10^6^ viable bacteria was administered transendoscopically, with 20 ml infused into the right mainstem bronchus and 20 ml into the left mainstem bronchus via a sterilized silastic tube inserted into the endoscope channel. The silastic tube was flushed twice with 20 ml of air after each 20-ml bacterial infusion. Foals and their mares were housed individually in stalls and separately from other mare and foal pairs for 1 week following experimental infection. After 1 week, these mare/foal pairs were transferred back to their original pasture.

### Quantitative PCR

Trizol reagent (Invitrogen) was used for total RNA extraction according to the manufacturer’s protocol. RNA was isolated using Direct-zol RNAeasy kits (Zymogen). cDNA was synthesized with BioRad iScript Direct Synthesis kits (BioRad) according to the manufacturer’s protocol. qRT-PCR was performed in triplicate wells using PowerUp SYBR Green Master Mix. Data were analyzed on a QuantStudio 6 Real-Time PCR System (Applied Biosystems).

### RNA-sequencing

RNA was isolated from cells in biological triplicate using Trizol reagent (Invitrogen) and Direct-zol RNAeasy miniprep kit (Zymogen) according to the manufacturer’s protocol. Agilent Technologies Bioanalyzer 2100 (Agilent Technology, Santa Clara, CA US) was used to verify RNA integrity number (RIN), rRNA ratio and RNA concentration. RNA-seq and library prep was performed by Texas A&M AgriLife Research Genomics and Bioinformatics Service. High-throughput RNA sequencing of samples was carried out on an Illumina NovaSeq 6000 S1 X using 2× 100-bp paired-end reads, which generated an average of ~32 million raw sequencing reads from each of 3 biological replicates for uninfected and infected macrophages. Analysis was performed as previously described (71). Briefly, raw reads were filtered and trimmed and Fastq data was mapped to the Mus musculus Reference genome (RefSeq) using CLC Genomics Workbench (Qiagen). Differential expression analyses were performed using CLC Genomics Workbench. Relative transcript expression was calculated by counting reads per kilobase of exon model per million mapped reads (RPKM). Statistical significance was determined using the EDGE test via CLC Genomics Workbench. The differentially expressed genes were selected as those with a p-value threshold <0.05.

### ISRE Reporter Assay

Macrophage-secreted type I IFN levels were determined using a L929 cells stably expressing a luciferase reporter gene under the regulation of type I IFN signaling pathway (L929 ISRE cells). At the indicated times post-infection, macrophage cell culture media was harvested and stored at −80°C. On the day prior to the assay, 5×10^4^ L929 ISRE cells were added to each well of a white 96-well flat-bottomed plate and incubated at 37°C/5%CO_2_ overnight. On the day of the bioassay, a 1:5 dilution of media from infected macrophages was added to each well of L929 ISRE cells, then incubated for 5h. Cells were washed with 1X PBS, lysed in reporter lysis buffer, then 30 μl of Luciferase Assay System solution (Promega) added and luminescence read immediately using a Cytation5 plate reader.

### IFN-β ELISA

Macrophage supernatant samples were harvested at 0-, 8-, and 24h post infection and stored at - 80°C until thawing on the day of the assay. Invitrogen Nunc MaxiSorp 96-well plates were coated with 50 μl of capture antibody diluted 1:5000 (Santa Cruz, sc-57201) in 0.1 M carbonate buffer and were incubated at 4°C overnight. Wells were then blocked using PBS+10% FBS for 2h at 37°C. Fifty μl of undiluted sample was added to each well. IFN-β standard (PBL, 12400-1) was diluted 1:4 for serial dilutions and incubated at room temperature (20°C) overnight. Samples were washed with PBS+0.05% TWEEN before each step. Detection antibody (RnD Systems, 32400-1) was added at a 1:2000 dilution and incubated at room temperature (20°C) overnight. After washing, secondary antibody (Cell signaling technology, 7074) was added to each well at a 1:2000 dilution and incubated for 3h. Following washing, the reaction was visually monitored until the standard was developed, then TMB substrate (SeraCare) was added and the reaction stopped with 2N H_2_SO_4_. The ELISA was read immediately at 450 nm using a BioTek plate reader.

### Western Blot

Cell monolayers were washed with 1X PBS and lysed in 1X RIPA buffer (150 mM NaCl, 1.0% NP-40, 0.5% sodium deoxycholate, 0.1% SDS, 50 mM Tris, pH 8.0) with protease and phosphatase inhibitors (1 tablet per 10 ml; Pierce). DNA was degraded using 250 units benzonase (EMD Milipore). Proteins were separated by SDS-PAGE and transferred to nitrocellulose membranes. Membranes were blocked for 1 h at RT in Odessey blocking buffer (Licor) or 4% BSA and incubated overnight at 4°C with the following antibodies: pNF-κB Ser536, 1:1000 (Cell Signaling 3033S), STAT1 1:1000 (Cell Signaling 9172S), pSTAT1 Tyr701 1:1000 (Cell Signaling 9177S); IRF3 1:1000 (Cell Signaling 4302), pIRF3 Ser396 1:1000 (Cell Signaling 4947); Beta Actin 1:5000 (Abcam 6276), Tubulin 1:5000 (Abcam 179513). Membranes were washed 3x in 1X TBS 0.1% Tween 20, and incubated with appropriate secondary antibody (Licor) for 2 h at RT (20°C) prior to imaging on Odessey Fc Dual-Mode Imaging System (Licor).

### Immunofluorescence microscopy

RAW 264.7 macrophages expressing epitope tagged galectins were seeded at 2×10^5^ cells/well on glass coverslips in 24-well dishes. At the indicated time point, cells were fixed in 4% paraformaldehyde for 10 min at 37°C, then washed three times with PBS. Coverslips were incubated in primary antibody diluted in TBS+ 0.25% Triton-X + 5% Normal Goat Serum (TBST-NGS) for 3h. Cells were then washed three times in PBS and incubated in secondary antibodies diluted in TBST-NGS for 1h. Coverslips were incubated in DAPI for 5 minutes, then washed twice with PBS and mounted on glass slides using Fluoromount (Diagnostic Biosystems; K024) for imaging. Images were obtained using a FV1200 Olympus inverted confocal microscope equipped with 60X oil immersion objective. Images were analyzed using ImageJ. Maximum intensity projections of z-stacks were obtained and projected images were thresholded such that Flag-tagged galectin puncta in macrophages were masked and counted. To obtain percent colocalization, GFP-tagged *R. equi* 103 were thresholded until they were masked, and region of interest (ROI) was saved. The *R. equi* ROI was then applied to the thresholded galectin channel and measured. Results are expressed as percentage of bacteria colocalized with galectin.

### Statistics

All data are representative of at least 2 independent experiments with n ≥ 3. Statistical analysis was performed using GraphPad Prism software (GraphPad, San Diego, CA). Two-tailed unpaired Student’s t-tests were used for statistical analysis unless otherwise noted. Results are reported as the mean ± SD.

### Data availability statement

The data that support the findings of this study are available from the corresponding author, ROW, upon reasonable request.

## ACKNOWLEDGEMENTS

We would like to thank members of the Patrick and Watson labs for their critical reading and feedback on this manuscript. We would like to acknowledge the Texas A&M AgriLife Research Genomics and Bioinformatics Service for performing our RNA-sequencing experiments. This work was supported by funds from the National Institutes of Health, grant 1R01AI125512 (ROW), the Texas A&M University T3 grant (ROW, KLP, AIB), the Link Equine Research Endowment at Texas A&M University (NDC), TAMU CVM Core Facility grant (KJV), and NIH training grant 5T32OD011083-10 (KJV). We gratefully acknowledge Dr. Mary Hondalus for provision of GFP expressing *R. equi* ATCC 103, and Kelsi West for Trif and MyD88 KD macrophage cell lines.

