## Supplemental Figure 1 for "The opportunistic intracellular bacterial pathogen *Rhodococcus equi* elicits type I interferons by engaging cytosolic DNA sensing in macrophages"

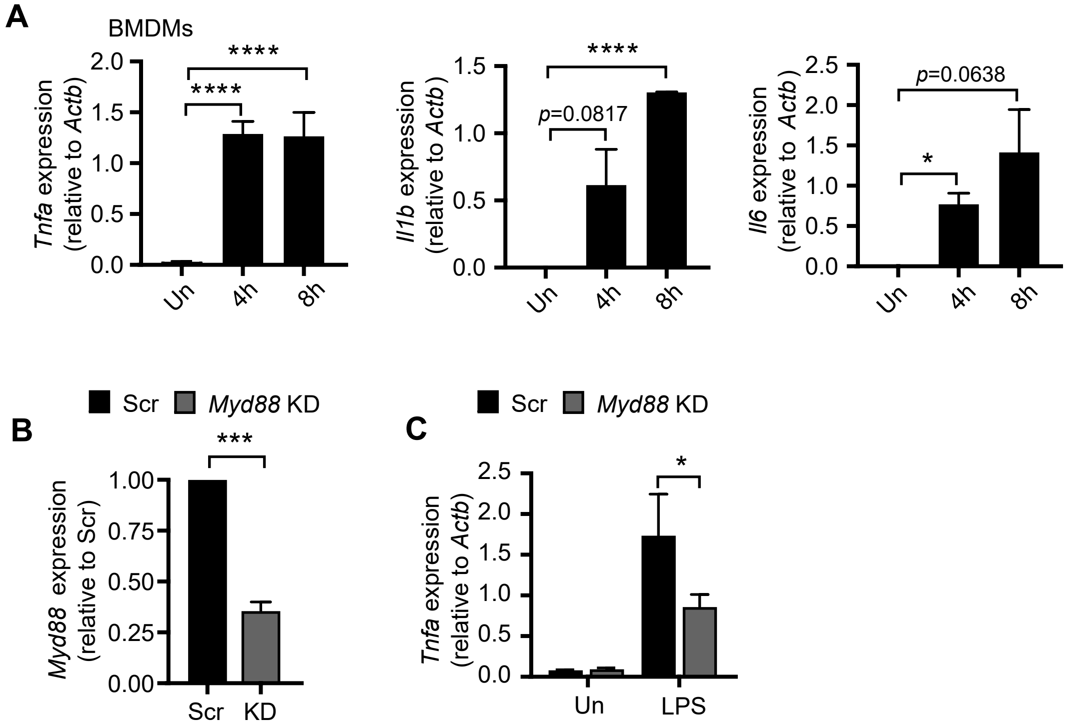


**Supplemental Figure 1** (A) qRT-PCR of Tnfa, Il1b and Il6 in BMDMs at 4 and 8 hours post infection with R. equi. (B) qRT-PCR of Myd88 in RAW 264.7 macrophages with Scr control. (C) qRT-PCR of Tnfa in MyD88 KD and Scr macrophages treated with LPS for 4 hours. RT-qPCR represent 3 biological replicates ± SD, n=3. For all experiments in this study, statistical significance was determined using Students' t-test. *p < 0.05, **p < 0.01, ***p < 0.001, ****p < 0.0001, n.s. = not significant.
