## Supplemental Figure 2 for "The opportunistic intracellular bacterial pathogen *Rhodococcus equi* elicits type I interferons by engaging cytosolic DNA sensing in macrophages"

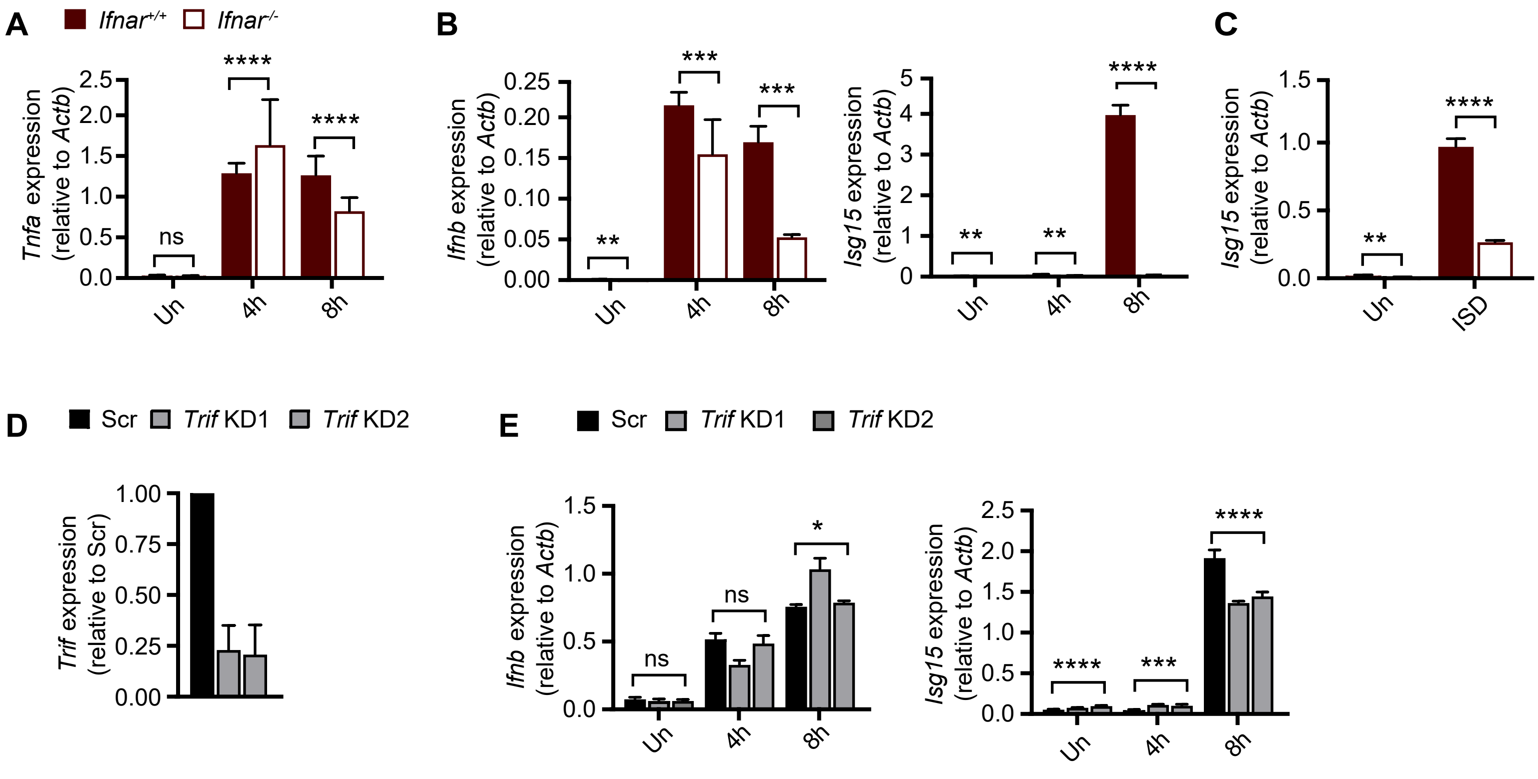


**Supplemental Figure 2** (A) RT-qPCR of Tnfa in IFNAR BMDMs infected with R. equi for the indicated times. (B) As in A but Ifnb and Isg15. (C) qRT-PCR of Isg15 in IFNAR+/+ (WT) and IFNAR-/- (KO) BMDMs treated or not with ISD for 4 hours. (D) qRT-PCR of Trif in RAW 264.7 KD cells relative to Scr control. (E) qRT-PCR of Ifnb and Isg15 in Trif KD and Scr RAW 264.7 macrophages. RT-qPCR represent 3 biological replicates ± SD, n=3. For all experiments in this study, statistical significance was determined using Students' t-test. *p < 0.05, **p < 0.01, ***p < 0.001, ****p < 0.0001, n.s. = not significant.
