## Supplemental Figure 3 for "The opportunistic intracellular bacterial pathogen *Rhodococcus equi* elicits type I interferons by engaging cytosolic DNA sensing in macrophages"

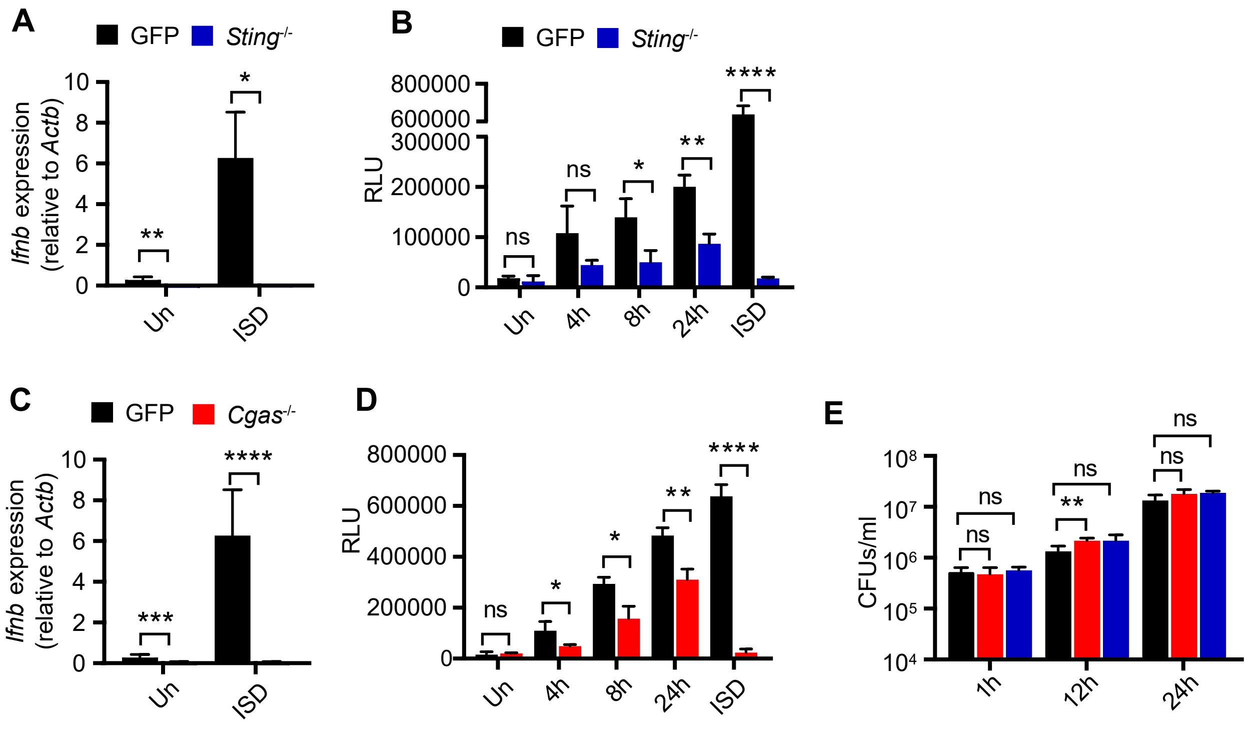


**Supplemental Figure 3** (A) RT-qPCR of Ifnb in GFP control and STING KO RAW 264.7 macrophages treated or not with ISD for 4 hours. (B) ISRE reporter assay measuring relative luminescence units as a readout for type I IFN protein levels secreted into supernatants from GFP control or STING KO RAW 264.7 macrophages infected or not with R. equi for the indicated times or treated with ISD for 4 hours. (C) As in F but in GFP control and cGAS KO RAW 264.7 macrophages. (D) As in G but in GFP control and cGAS KO RAW 264.7 macrophages. (E) CFUs of GFP control, cGAS KO and STING KO RAW 264.7 macrophages infected with R. equi for the indicated times. RT-qPCR represent 3 biological replicates ± SD, n=3. For all experiments in this study, statistical significance was determined using Students' t-test. *p < 0.05, **p < 0.01, ***p < 0.001, ****p < 0.0001, n.s. = not significant.
